# GametesOmics: A Comprehensive Multi-Omics Database for Exploring the Gametogenesis in Humans and Mice

**DOI:** 10.1101/2023.09.05.556316

**Authors:** Jianting An, Jing Wang, Siming Kong, Shi Song, Wei Chen, Peng Yuan, Qilong He, Yidong Chen, Ye Li, Yi Yang, Wei Wang, Rong Li, Liying Yan, Zhiqiang Yan, Jie Qiao

**Affiliations:** State Key Laboratory of Female Fertility Promotion, Center for Reproductive Medicine, Department of Obstetrics and Gynecology, Peking University Third Hospital, Beijing 100191, China; National Clinical Research Center for Obstetrics and Gynecology (Peking University Third Hospital), Beijing 100191, China; MOE Key Laboratory of Assisted Reproduction (Peking University), Beijing 100191, China; Beijing Key Laboratory of Reproductive Endocrinology and Assisted Reproductive Technology, Beijing 100191, China; Peking-Tsinghua Center for Life Sciences, Peking University, Beijing 100871, China; Beijing Advanced Innovation Center for Genomics, Beijing 100191, China

**Keywords:** Gametogenesis, Oogenesis, Spermatogenesis, Transcriptomics, Epigenomics

## Abstract

Gametogenesis plays an important role in the reproduction and evolution of species. The transcriptomic and epigenetic alterations in this process can influence the reproductive capacity, fertilization, and embryonic development. The rapidly increasing single-cell studies have provided valuable multi-omics resources. However, data from different layers and sequencing platforms have not been uniformed and integrated, which greatly limits their use for exploring the molecular mechanisms that underlie oogenesis and spermatogenesis. Here, we developed GametesOmics, a comprehensive database that integrated the data of gene expression, DNA methylation, and chromatin accessibility during oogenesis and spermatogenesis in humans and mice. GametesOmics provides a user-friendly website and various tools, including *Search* and *Advanced Search* for querying the expression and epigenetic modification of each gene; *Analysis Tools* with *Differentially Expressed Gene (DEG) analysis* for identifying DEGs, *Correlation analysis* for demonstrating the genetic and epigenetic changes, *Visualization* for displaying single-cell cluster and screening marker genes as well as master transcription factors (TFs), and *MethylView* for studying the genomic distribution of epigenetic modifications. GametesOmics also provides *Genome Browser* and *Orthologs* for tracking and comparing gene expression, DNA methylations, as well as chromatin accessibilities between humans and mice. GametesOmics offers a comprehensive resource for biologists and clinicians to decipher the cell fate transition in germ cell development, and can be accessed at http://gametesomics.cn/.

## Introduction

Gametogenesis is a key process in reproduction and evolution of species, during which diploid germ cells undergo mitotic and meiotic divisions, then differentiate into mature haploid gametes including eggs (oogenesis) and sperms (spermatogenesis) [1,2]. The complex molecular regulatory mechanism involving genetic and epigenetic changes supports the successful genesis of the gametes [3-5]. Nowadays, the rapid development of single-cell sequencing technologies has broadened our understanding of this process and provided a large volume of omics data [6-10]. However, these data have not been integrated or normalized for utilization, which hampers biologists and clinicians to explore the molecular mechanisms that underlie oogenesis as well as spermatogenesis.

Several databases contain valuable omics data of human and mouse. DISCO database collects the single-cell RNA-seq datasets covering various tissues and cell types in human, which provides a comprehensive platform for exploring gene expressions across different cell types and human tissues [11]. scMethBank database includes DNA methylation data across a several species including human and mouse and provides many useful functions such as search, browsing, and visualization for researchers [12]. dbEmbryo and DevOmics integrate multi-omics sequencing data across different developmental stages in human and mouse early embryos [13,14]. EmAtlas exhibits the spatiotemporal landscape of human and mouse embryos from the perspective of cell, tissue, genome, gene and protein levels [15]. These current databases greatly promote the understanding of the genetic and epigenetic mechanisms in the development of human and mouse. However, they may lack the collection of comprehensive multi-omics sequencing data of the gametogenesis and online tools for exploring specific stepwise process regarding the oogenesis and spermatogenesis, which may hinder the exploration into the landscape and regulatory mechanisms of the development process during oogenesis and spermatogenesis. Therefore, it is highly desirable to construct a customized database that includes multi-omics data across complete developmental stages and offers user-friendly tools for identifying key regulators of cell fate transition in gametogenesis.

With the aim of establishing a comprehensive unified atlas of oogenesis and spermatogenesis, we developed GametesOmics (http://gametesomics.cn/), a multi-omics database which integrated the data of gene expression, DNA methylation and chromatin accessibility spanning non-growing oocyte, growing oocyte, fully-grown oocyte, metaphase Ⅰ oocyte as well as metaphase Ⅱ oocyte during oogenesis, and spermatogonia stem cell, spermatogonia, spermatocyte, spermatid as well as mature sperm during spermatogenesis in human and mouse. It provides various functions for querying and displaying the gene expression as well as epigenetic modification changes, performing differential analysis, visualizing single-cell clusters, identifying master TFs and tracking homologous genes’ expressions in human and mouse. GametesOmics helps researchers study the synergistic change between different layers and explore human and mouse homologous key factors that trigger the development of the gametes, so as to decode regulatory mechanisms in cell fate determination in gametogenesis.

### Database implementation

#### Data collection

We retrieved studies in PubMed (https://pubmed.ncbi.nlm.nih.gov/), Gene Expression Omnibus (GEO, https://www.ncbi.nlm.nih.gov/geo/), Sequence Read Archive (SRA, https://www.ncbi.nlm.nih.gov/sra/), as well as Genome Sequence European Nucleotide Archive (ENA, https://www.ebi.ac.uk/ena/browser/) using the arrangement and combination of the keywords including “gametogenesis”, “oogenesis”, “oocyte”, “spermatogonia”, “sperm”, “RNA-seq”, “transcription”, “methylation”, “chromatin accessibility”, and obtained related studies containing the single-cell sequencing datasets of gametogenesis in human and mouse. These studies used appropriate number of samples to demonstrate their discoveries. In consideration of the data quality and comparability, we made efforts to choose the datasets that cover the most complete developmental stages, with the largest number of samples and the most advanced sequencing methods as well as the same sequencing platforms to ensure the data consistency as possible.

We collected 6,689 samples generated by RNA-seq for gene expression (4,381 samples), whole-genome bisulfite sequencing / COOL-seq (WCG) for DNA methylation (1,162 samples) and COOL-Seq (GCH) for chromatin accessibility (1,146 samples) across oogenesis (7 stages for human and mouse) and spermatogenesis (14 stages for human and 20 stages for mouse) [6-10]. We downloaded the raw data in FASTQ form and processed them with our unified pipeline for raising their comparability as described below.

Additionally, we added gene expression datasets (10 samples for human oogenesis, 14 samples for mouse oogenesis, 411 samples for monkey oogenesis; 10,115 samples for human spermatogenesis, 29,552 samples for mouse spermatogenesis, 19,476 samples for monkey spermatogenesis) from other data sources [16-19] as supplement and integrated them into *Advanced Search*.

#### Processing of scRNA-seq data

First, the reads of pooling library were separated into single cell by the barcodes as previously described [9]. The reads of each cell were filtered and trimmed with Trim Galore! (v0.6.6) using default parameters. Next, the trimmed reads were aligned to hg38 (human) or mm10 (mouse) reference genome using STAR (v2.7.1a) with default parameters. The rRNA reads were removed by RSeQC (v2.3.7) with default parameters. Only uniquely mapped reads were retained and the duplications were removed based on UMI information. The Reads of exon model per million mapped reads (RPM) of each gene were calculated to represent the gene expression.

For the visualization of gene expression features, we used Seurat (v4.0) package to perform the RunTSNE function and RunUMAP function (dims=1:20) based on normalized RPM expression values. The marker genes of each developmental stage were identified by FindAllMarkers function (min.pct = 0.25, logfc.threshold = 0.5, return.thresh = 0.01). Then we used the homer software to perform the motif enrichment to find the TFs regulating the stage-specific marker genes with default parameters.

For the exhibition of gene expression in the *Genome Browser*, BAM files in same developmental stage were merged and transferred into BIGWIG form with 50 bp windows using bamCoverage function in deepTools (v3.4.3).

#### Processing of scCOOL-seq data and bisulfite sequencing data

The raw data were filtered and trimmed with Trim Galore! and then aligned to hg38 (human) or mm10 (mouse) reference genome with Bismark (v0.22.3) in single-end mode. The PCR duplicates were removed by samtools. We used bismark_methylation_extractor function of Bismark to calculate methylation level or chromatin accessibility of each covered cytosine. The number of “methylated” reads (reported as C) was divided by the total number of “methylated” and “unmethylated” reads (reported as C or T) at the same reference position. The average DNA methylation level was represented by the average level of WCG sites (including ACG or TCG) and chromatin accessibility was represented by the average level of GCH sites (including GCA, GCC or GCT). The methylation level and chromatin accessibility in gene promoters were evaluated by the average WCG sites or GCH sites within the regions from 1.5 kb upstream to 0.5 kb downstream of transcription start site for each gene.

#### Database interface

We collected 66,267 samples (6689 samples for primary datasets and 59,578 samples for additional datasets) from different developmental stages in oogenesis (mainly divided into non-growing oocyte, growing oocyte, fully-grown oocyte, metaphase Ⅰ oocyte, and metaphase Ⅱ oocyte) and spermatogenesis (mainly divided into spermatogonia stem cell, spermatogonia, spermatocyte, spermatid, and mature sperm) [6-10,16-19] (Tables S1 and S2; **Figure 1**A). Then we processed the data with our unified pipeline and developed several tools for studying synergistic changes of different layers in gametogenesis (Figure 1B).

**Figure 1.**
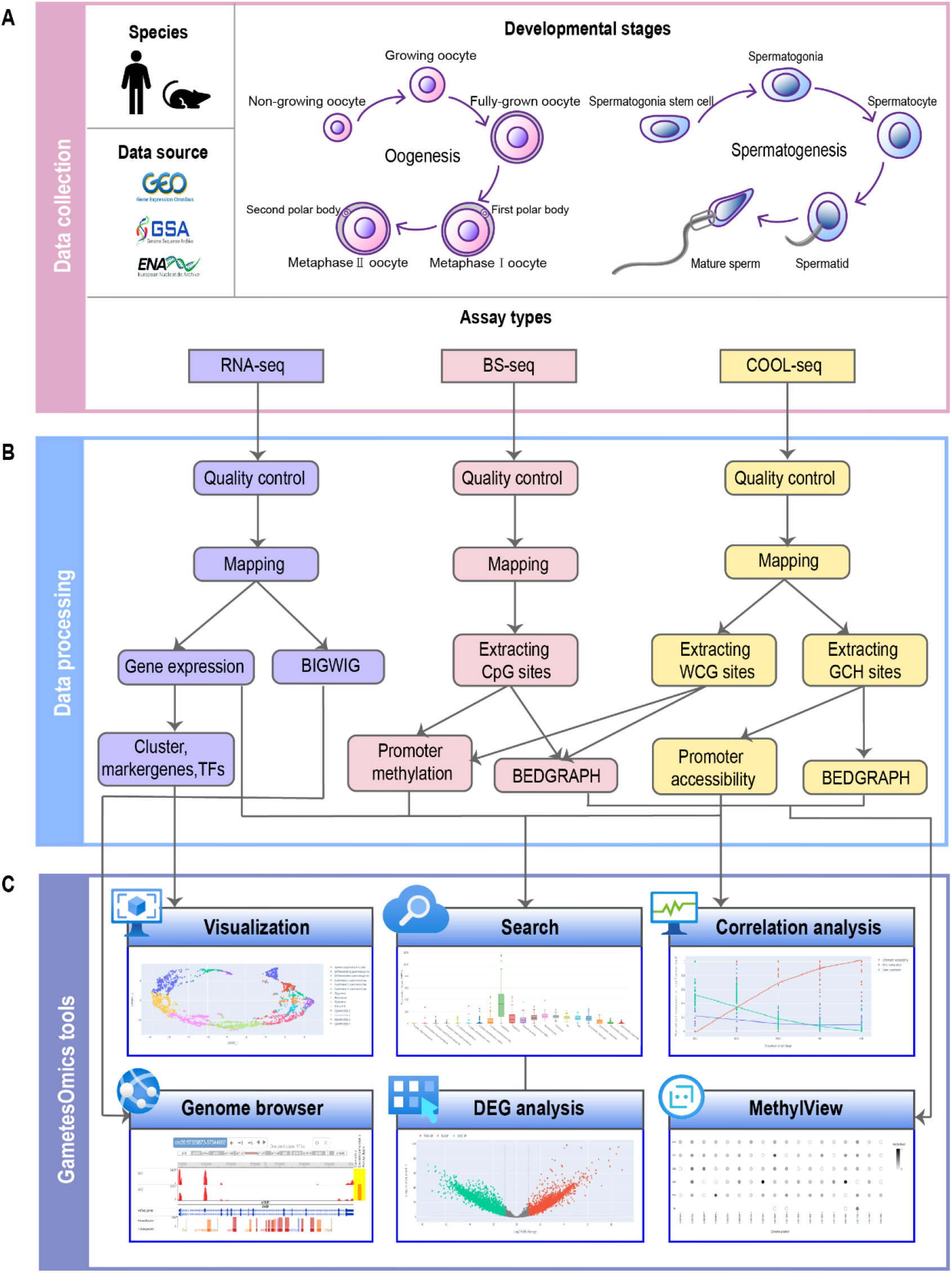
The framework of GametesOmics. **A**. Data collection of multi-omics sequencing data. **B**. The custom pipeline to process the collected data for downstream applications and display in GametesOmics. **C**. The tools provided by GametesOmics.

GametesOmics provides a user-friendly website as well as various useful functions including: 1) *Search* and *Advanced Search* for querying and displaying expression and epigenetic modification levels of interested genes; 2) *Analysis Tools* with *DEGs analysis* for applying differential analysis, *Correlation analysis* for simultaneously displaying the dynamic changes in gene expression and coordinated epigenetic alterations, *Visualization* for showing marker genes as well as master TFs, and *MethylView* for viewing the methylation in WCG sites or GCH sites; 3) *Genome Browser* and *Orthologs* for tracking and comparing expressions and modifications of homologous genes between human and mouse (Figure 1C).

#### *Search* and *Advanced Search*

The primary goal of *Search* and *Advanced Search* is to help users to retrieve the expression and epigenetic modification changes of their interested genes.

Users can input a gene (either gene symbol, Entrez ID or Ensembl ID) then choose human or mouse in *Search* box on the home page. They will get quick access to the result page of their queried gene, which includes: the detailed descriptions (such as the gene official name, alias, functions, related gene ontology term and Reactome pathway); gene expression levels, promoter methylation levels as well as promoter chromatin accessibility across the developmental stages in oogenesis or spermatogenesis.

*Advanced Search* supports users to input multiple genes at one time to access their expressions or modifications in promoters. Users can also choose different datasets that they are interested in. This function may not only save the time when searching large batches of genes, but also display the correlation of these genes with various plot forms (such as line plot, bar plot, box plot, or heatmap).

### Analysis tools

#### DEGs analysis

DEGs between two developmental stages usually participate in driving the changes in the morphology or functions during the germ cell development. Therefore, identifying DEGs is important for screening the key genes which have the potential to determine the fate of oocytes or sperms.

By choosing the species, gametes (oocyte or sperm), assay types, developmental stages in the option page, the DEGs can be obtained by limma-voom method [20] between the selected stages. The result page will show the clusters of samples, heatmap of top 20 DEGs, as well as the DEG-enriched gene ontology terms and Reactome pathways.

#### Correlation analysis

It has been revealed that the dynamic changes of epigenetic alterations including DNA methylation and chromatin accessibility can regulate gene expression in gametogenesis. Studies found that a number of gene promoters were accessible in the human growing oocytes at an early stage to prepare for their transcription [9,21]. Therefore, the coordinate changes and correlation between the gene expression and epigenetic modifications may help us to delineate the regulatory mechanism in gametogenesis.

*Correlation analysis* provides a tool for users to view and compare the coordinate changes in expression levels, methylation levels as well as chromatin accessibility of their interested genes across different developmental stages in gametogenesis. Researchers may be able to explore the relationship between the gene expression and epigenetic modifications, and to decode the molecular mechanism of the germ development and inheritance.

#### Visualization

Gametogenesis is a highly ordered and continuous process, which can be reflected and traced by the single cell gene expression. Besides, by exploring the gene expression characteristics along the developmental trajectory, the biomarkers and key TFs of each stage could be screened.

*Visualization* is used to display the gene expression features of each single cell. Users can select species, gamete, and cluster method in the option page. Then the result page will provide a visualization of the single-cell RNA-seq data. First, clustered plot exhibits the gene expression similarities and distance between these single cells. The developmental stages are marked with different colors, which may show the developmental trajectories of oogenesis or spermatogenesis. Second, users can obtain the heatmap of top 20 marker genes of any developmental stage. Third, upstream TFs regulating all the marker genes of this developmental stage would be displayed with bubble plot. Using this module, users may be able to find the master TFs with high expression levels as well as critical factors and explore the regulatory mechanisms of the development process.

#### MethylView

The epigenetic modifications in different genomic regions can cause different regulatory effects on the gene expression. For instance, it was revealed that hypermethylation in promoters or enhancers usually resulted in transcriptional silencing, while hypomethylation in gene bodies usually promoted the gene expression [22]. Thus, accessing the epigenetic modifications across the different genomic regions could help us discuss their regulation effects of transcription more specifically.

*MethylView* provides an access to the lollipop diagrams of each WCG site (reflecting the DNA methylation level) or GCH site (reflecting the chromatin accessibilities) of selected species, gametes and genomic regions. The color of each dot shows the level of DNA methylation or chromatin accessibilities. Using this tool, users will directly view the epigenetic modification levels in any region such as intron, exon, intergenic region and so on, which would make the investigation of the epigenetic regulation more easily.

#### *Genome Browser* and *Orthologs*

*Genome Browser* provides a window for users to directly view gene expression and epigenetic modification tracks of their interested developmental stages in a specific gene location or chromosome region. This function exhibits the transcription and modification on the reference genome in detail.

Mouse are usually used as the mammalian experimental model in genetic studies for their similar genome with human, easy manipulation on the genome, and short lifecycle [23]. Therefore, we can utilize result of the gene research in mouse as the reference for human studies according to the corresponding genomic homologous regions. *Orthologs* was developed to view the expression and modification of any human-mouse homologous regions in the selected developmental stage. This function will not only help researchers to study the similarities as well as the divergences between human and mouse, but will also promote the transformation from the experimental researches into clinical applications.

### Applications

To present the utilization of GametesOmics, we use it for exploring the essential genes in the developmental stages of oogenesis and spermatogenesis. We applied the *Visualization, Search, Advanced Search, DEGs analysis*, and *Ortholog* tools for the case studies.

### Screening the regulatory factors of spermatids in human and mouse

Spermatogenesis process mainly includes the spermatogonia proliferation and differentiation, the spermatocyte meiosis and spermatid post-meiotic development [24]. The post-meiotic development, also called spermiogenesis, involves the formation of acrosome and flagella, condensation of the chromatin, elimination of extra cytoplasm, and transforming into spermatozoa [25,26]. This process is highly orchestrated and depends on the accurate regulation of many genes. However, the reports of the key factors of the spermatid post-meiotic development are limited.

In mouse, the single-cell spermatids displayed by U-map plots are at the end of the spermatogenesis trajectory (Figure 2A). Among the spermatids, “spermatids.steps3to4” sub-stage was revealed to be quite essential as the single sphere acrosomal granule starts to form and the genes related to chromatoid body formation, cilium movement, as well as acrosome formation also reached their peak levels [8,27]. We utilized the *Visualization* function in GametesOmics to investigate the master TFs of this developmental stage. The bubble plot shows that the promoters of stage-specific genes are enriched in *Rfx2* motif and *Rfx2* is highly expressed in spermatids (Figure 2B, 2C). Previous studies have found that the spermiogenesis in male *Rfx2*^− /−^ mice was disrupted. Round spermatids failed to generate flagella and were not able to differentiate into elongated spermatids, along with altered expressions of a large number of genes associated with the spermatogenesis, and eventually underwent apoptosis [28]. RNA-Seq and ChIP-Seq analysis have identified genes directly controlled by *Rfx2* during spermiogenesis and found that they were involved in cilium function, cytoskeleton remodeling and cell adhesion, which indicated that *Rfx2* is an important TF for the round spermatid differentiation as well as flagellar biogenesis [29]. This verified the analytical result of the *Visualization* function.

**Figure 2.**
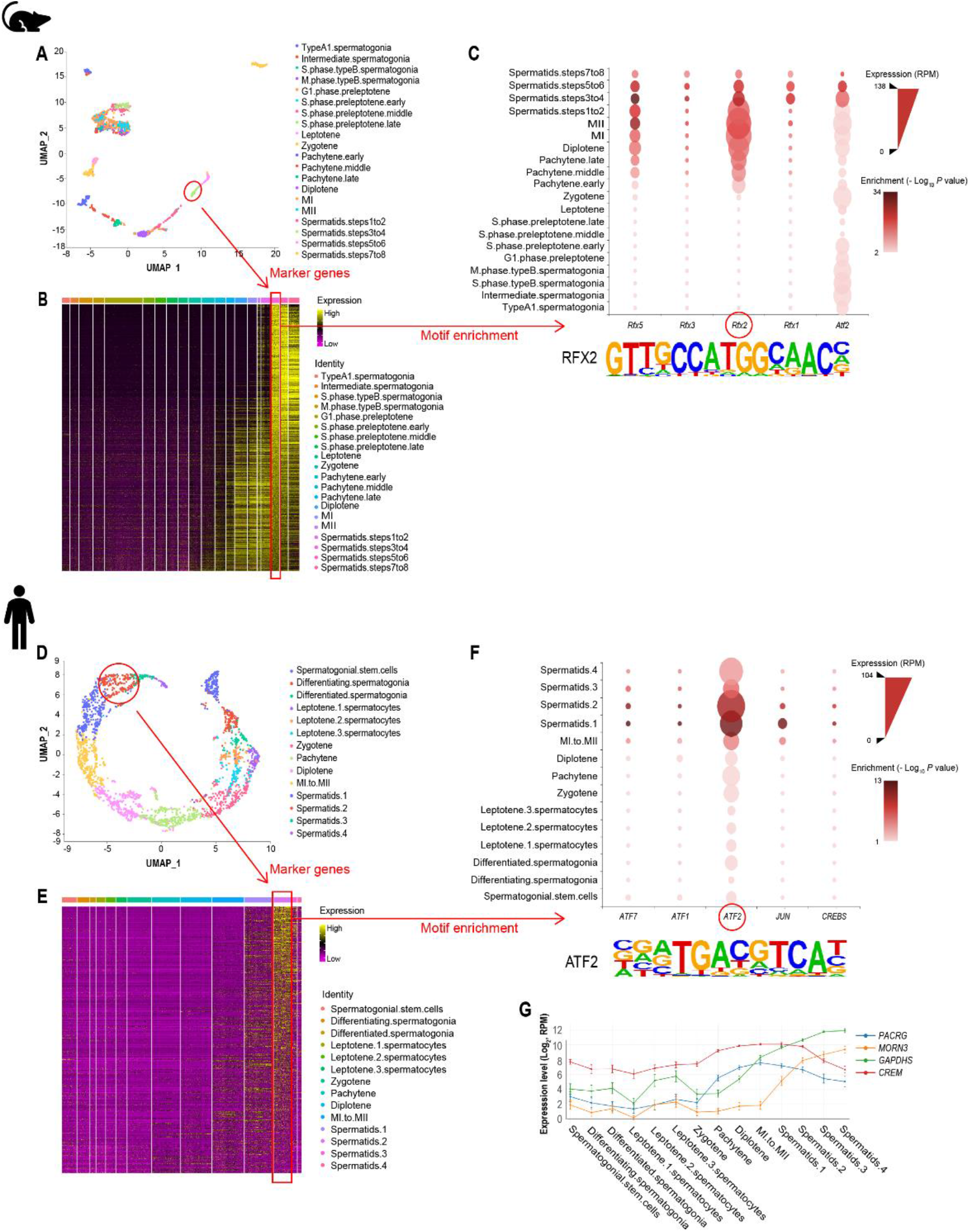
Screening the regulatory factors of spermatids. **A**. U-map plots showing the mouse spermatogenesis trajectory. Developmental stages are marked by different colors. “spermatids.steps3to4” stage is highlighted. **B**. Heatmap showing the stage-specific genes of “spermatids.steps3to4” stage in mouse spermatogenesis. **C**. The bubble plots of top5 transcription factors (TFs) enriched in the promoters of stage-specific genes of “spermatids.steps3to4” stage, with selected RFX2 motif displayed below. **D**. U-map plots showing the human spermatogenesis trajectory. Developmental stages are marked by different colors. “spermatids.2” stage is highlighted. **E**. Heatmap showing the stage-specific genes of “spermatids.2” stage in human spermatogenesis. **F**. The bubble plots of top5 TFs enriched in the promoters of stage-specific genes of “spermatids.2” stage, with selected ATF2 motif displayed below. **G**. Line plot showing the expression of *PACRG, MORN3, GAPDHS*, and *CREM* during human spermatogenesis.

In human, the single-cell spermatids are also at the end of the spermatogenesis trajectory (Figure 2D). Among the spermatids, “spermatids.2” sub-stage is important, in which the peanut agglutinin, a sperm-acrosome-specific marker, shapes as crescent and the transition nuclear proteins switches to protamine at this stage [8]. In the same way, we tend to find the key TF in human using the *Visualization* function. We found that *ATF2* is highly expressed and enriched in human spermatids (Figure 2E, 2F), which have not been studied in human spermatogenesis. *ATF2* has been reported to play an important role during human early development. It can bind the promoters of many key genes regulating the inflammatory signaling, cell cycle control, and glycosylation et al [30-32].

We downloaded the target genes of *ATF2* from Cistrome database (http://cistrome.org/db/). The downstream includes several regulatory genes that related to the development of the sperm, such as *PACRG, MORN3, GAPDHS*, and *CREM*. We input these downstream target genes above into the *Advanced Search* tool, it turned out that they were highly expressed in the spermatids in human (Figure 2G). *PACRG* is found to be highly expressed in mouse testis and is required to build the sperm flagella [33]. And loss of *PACRG* can cause male sterility [34]. In human, null mutations in *PACRG* is observed in patients with severe sperm motility disorders [35]. *MORN3* is highly expressed in male germ cells of mouse and is localized in the acrosome and manchette, which are also specific and essential structures for spermiogenesis [36]. *GAPDHS* is a sperm-specific expressed gene and play an essential role in energy production for sperm motility and male fertility. It has been also studied as a potential male contraceptive target for human [37]. *CREM* is reported to be important in the structuring of mature spermatozoa [38]. Lack of the *CREM* gene can cause mouse becoming infertile and the arrest of the round spermatids. Besides, it has been confirmed to be expressed in human germ cells and might act as a switch for the human spermatogenesis [39]. In conclusion, it indicated that the screened *ATF2* by GametesOmics might be a key regulator of the spermatid post-meiotic development in human.

### Screening the key genes of the growth of oocytes in human and mouse

In mammalian, the small and non-growing oocytes need weeks or months to grow into more than 100-fold of their volume [40]. In this process, they undergo a burst of transcription and translation to reach their full size, which is regarded as one of the foundations for fertilization and embryogenesis [41]. Growing oocyte I (GO1) is the main stage characterized by the rapid growth compared with fully-grown oocyte (FGO). Therefore, genes highly expressed in GO1 are essential for oogenesis in human and mouse. However, the cross-species conservation and diversity of gene expressions as well as epigenetic modifications between human and mouse need to be further investigated.

To explore the key regulatory genes of the growth of oocytes in human and mouse, we use *DEGs analysis* in GametesOmics for obtaining the DEGs between GO1 and FGO (Figure 3A). It turned out that 4,845 genes for human and 733 genes for mouse are relatively highly expressed in GO1 (Figure 3B). GO analysis indicated the unique human DEGs enriched in terms such as “protein transport”, “protein folding” and “ribosome biogenesis”, while the unique mouse DEGs enriched in terms such as “multicellular organism development”, “negative regulation of apoptotic process” and “chromosome organization” (Figure 3C, 3D). The 143 human and mouse overlapping genes are enriched in “positive regulation of transcription”, “proliferation”, as well as “growth factor signaling”, which implied that they might be conservatively expressed across species and function in the growing oocyte in human and mouse (Figure 3E).

**Figure 3.**
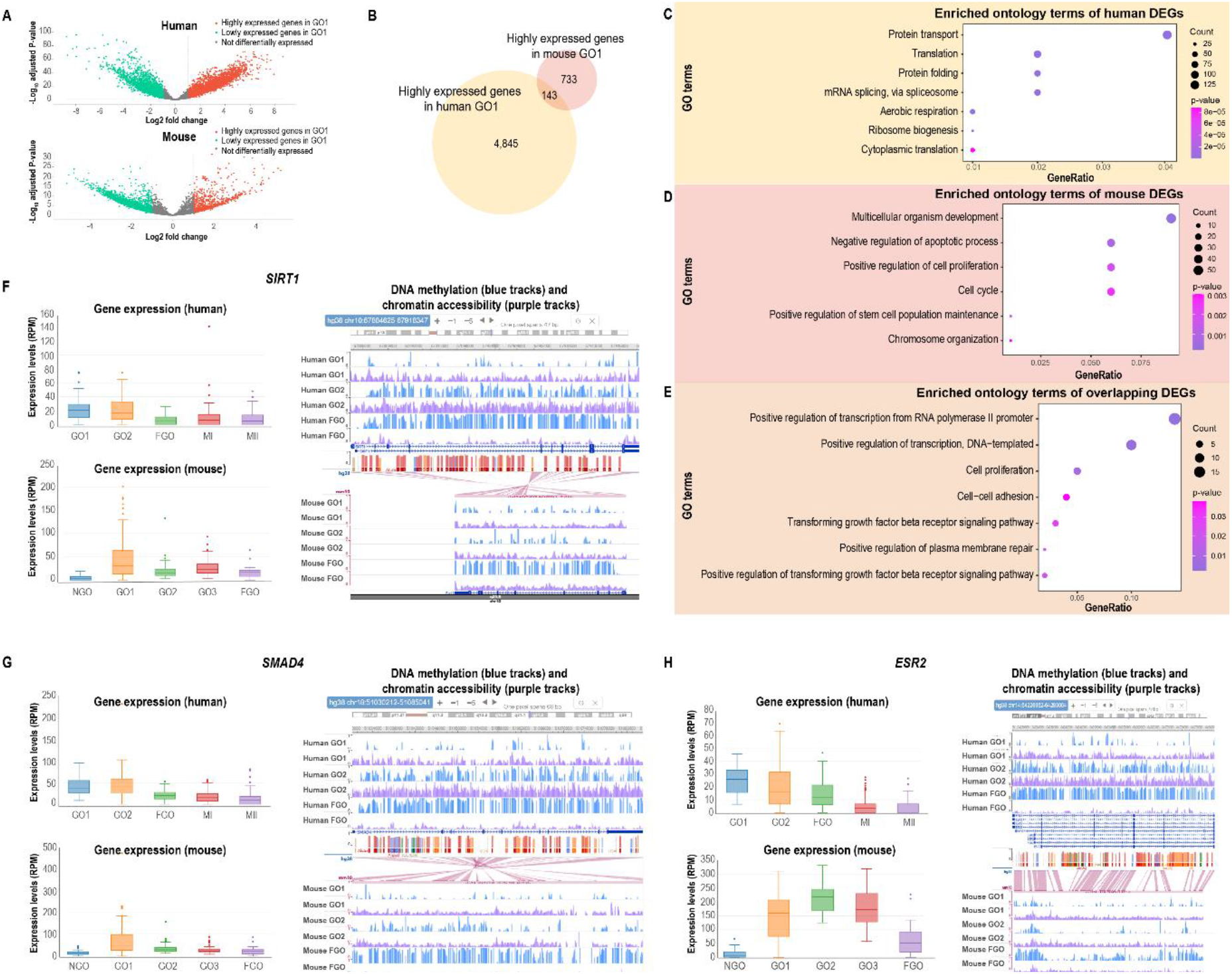
Screening the key genes of oocyte growth in human and mouse. **A**. The volcano plot showing the foldchange and P-value of differentially expressed genes (DEGs) between growing oocyte I (GO1) and fully-grown oocyte (FGO). **B**. The Venn diagram of highly-expressed genes in human and mouse GO1 stage. **C**. Gene ontology enrichment of the genes that are highly-expressed in human GO1. **D**. Gene ontology enrichment of the genes that are highly-expressed in mouse GO1. **E**. Gene ontology enrichment of the overlapping genes that are highly-expressed in human and mouse GO1. **F**. The gene expression levels (showed by the box plot), the DNA methylation tracks (showed by UCSC genome browser in blue) as well as chromatin accessibility tracks (showed by UCSC genome browser in purple) of *SIRT1/Sirt1* in human and mouse oogenesis. **G**. The gene expression levels (showed by the box plot), the DNA methylation tracks (showed by UCSC genome browser in blue) as well as chromatin accessibility tracks (showed by UCSC genome browser in purple) of *SMAD4*/*Smad4* in human and mouse oogenesis. **H**. The gene expression levels (showed by the box plot), the DNA methylation tracks (showed by UCSC genome browser in blue) as well as chromatin accessibility tracks (showed by UCSC genome browser in purple) of *ESR2*/*Esr2* in human and mouse oogenesis.

Then we also found that these overlapping genes could be regulated by divergent molecular mechanisms. For example, Sirtuin 1 (SIRT1) is one of the family of NAD-dependent deacetylases and is associated with metabolic challenges, DNA repair, stress response, and reproductive aging [42,43]. It also participates in chromatin organization by remodeling chromatin structure and accessibility. In mouse, the aging oocyte showed defective expression of SIRT1 protein [44]. In human, *SIRT1* has been proved to be related to the process of proliferation and activation of steroidogenesis [45]. *SIRT1* is highly expressed in the growing oocyte (Figure 3F). As shown in the result of *Ortholog* in GametesOmics, as for human, the low DNA methylation and high chromatin accessibility levels of promoter might promote the expression of *SIRT1* at GO1 stage. However, as for mouse, only the DNA methylation alterations were investigated to be associated with the regulation of *SIRT1* expression, while the chromatin accessibility showed relatively stable during the oogenesis (Figure 3F). SMAD4, which forms a complex with the phosphorylated R-SMADs, can regulate the gene for controlling the cell cycle and cell fate determination [45]. SMAD4 also functions as the central molecule of the TGF-β signaling pathway, which is associated with primordial follicle formation, ovulation, and signaling between the pituitary and ovary [46-48]. *SMAD4* was found to be specially expressed in oocytes and the follicular somatic cells. *Smad4* conditional knockout mice are observed to have reduced antral follicles, ovulation rates, as well as defects in cumulus cells. Their fertility decreased, and half are infertile by six months of age [49]. We found that the *SMAD4* is highly expressed in growing oocyte in the orthologs of human and mouse (Figure 3G), which implied that *SMAD4* might play an essential role in the oocyte growth in human as in mouse. Like *SIRT1, SMAD4* promoter also shows relatively low DNA methylation and high chromatin accessibility at this stage in human, while chromatin accessibility of SMAD4 is stable in mouse (Figure 3G). Besides, estrogen receptor 2 (ESR2) is an essential estrogen receptor and its polymorphisms and mutations are found to be associated with ovulatory dysfunctions in human [50,51]. As for mouse, *Esr2* loss of function can lead to defective follicle development and ovulation [52]. *ESR2* is highly expressed in human and mouse growing oocytes (Figure 3H) and showed a similar modification alteration with *SMAD4*.

In summary, the homologous DEGs identified by GametesOmics may play important roles in the growth of oocytes. Furthermore, the conservation and diversity in gene expression as well as epigenetic modification across human and mouse during gametogenesis can be explored through GametesOmics.

Collectively, GametesOmics provides informative datasets and useful tools for biologists to screen master TFs and regulatory genes during gametogenesis.

## Discussion

To our knowledge, GametesOmics is the first database which specifically deposited the multi-omics information of oogenesis and spermatogenesis in human and mouse. These single-cell sequencing data were processed with a unified pipeline, which made it viable for the quantization and comparation between these datasets. It provides various useful functions for searching and viewing gene expression and epigenetic modification, as well as tools for DEG identification, synergistic regulatory network investigation and master factor screening during gametogenesis. Users can easily utilize GametesOmics to analyze these integrated sequencing data according to their research demand.

Previous studies have portrayed genetic and epigenetic landscape in oogenesis as well as spermatogenesis, but the key factors driving the changes of morphology and function during these processes need to be further discovered. In our applications based on GametesOmics, we screened *Rfx2* as a potential master TF in mouse spermatids. Studies demonstrated that deletion of *Rfx2* disrupted the differentiation into elongated spermatids, which proved the practicability and reliability of our database. Also, based on our developed tools, we identified *SIRT1, SMAD4* as well a*s ESR2* as key genes in human and mouse oocyte growth. *SIRT1* was reported to be associated with proliferation metabolic challenges, DNA repair, and reproductive aging. *Smad4* conditional knockout reduced antral follicles and ovulation rates. *ESR2* encodes an essential estrogen receptor and is reported to be associated with follicle development and ovulation. This suggests the validity of these tools. We also found their conserved expressions and divergent epigenetic modifications in human and mouse, which supported the practicability of GametesOmics.

Taken together, we believe that GametesOmics will help researchers to dissect the molecular regulatory mechanisms in germ cell development from multilayers and facilitate their exploration of genetic and epigenetic inheritance between generations. Besides, it will also be possible to provide clues in deciphering and curing reproductive disorders as well as genetic diseases in clinic.

In the future, GametesOmics will continuously update and integrate more genetic and epigenetic sequencing data of single-cells across the oogenesis and spermatogenesis from newly published researches. Besides, we design to optimize and expand the functions according to the feedback of our users for offering more convenient applications as much as possible.

## Data availability

The GametesOmics database is available at http://gametesomics.cn/ and can be accessed without registration or login.

## CRediT author statement

**Jianting An**: Data curation, Methodology, Validation, Formal analysis, Investigation, Writing - original draft. **Jing Wang**: Data curation, Validation, Investigation. **Siming Kong**: Data curation, Resources. **Shi Song**: Resources, Investigation. **Wei Chen:** Resources, Investigation. **Peng Yuan**: Resources. **Qilong He**: Resources. **Yidong Chen**: Resources. **Ye Li**: Resources. **Yi Yang**: Resources. **Wei Wang**: Resources. **Rong Li**: Resources. **Liying Yan**: Conceptualization, Resources, Writing - review & editing, Supervision, Project administration. **Zhiqiang Yan**: Conceptualization, Resources, Methodology, Writing - review & editing, Supervision, Project administration. **Jie Qiao**: Conceptualization, Resources, Writing - review & editing, Supervision, Project administration, Funding acquisition. All authors approved the final version of the manuscript.

## Acknowledgments

This project is supported by the National Natural Science Foundation of China (Grant No. 82125013 and 82101741), the National Key R&D Program of China (2019YFA0801400) and Beijing Natural Science Foundation (7232203). The authors thank Prof. Fan Guo for providing detailed information about their previous study and Drs. Tong Chen as well as Pu Xue (EHBIO gene technology) for their technical assistance.

